# Reproducible Boolean model analyses and simulations with the CoLoMoTo software suite: a tutorial

**DOI:** 10.1101/2025.01.26.634971

**Authors:** Vincent Noël, Aurélien Naldi, Laurence Calzone, Loic Paulevé, Denis Thieffry

## Abstract

This tutorial provides stepwise instructions to install the 20 tools integrated in the CoLoMoTo software suite, to develop reproducible dynamical analyses of logical models of complex biological molecular networks.

The tutorial specifically focuses on the analysis of a previously published model of the regulatory network controlling mammalian cell proliferation. It includes chunks of python code to reproduce several of the results and figures published in the original article and further extend these results with the help of a selection of tools included in the CoLoMoTo suite.

The notebook covers the visualisation of the network with the tool GINsim, an attractor analysis with bioLQM, the computation of synchronous attractors with BNS, the extraction of modules from the full model, MaBoSS simulations of the wild-type model, as well as of selected mutants, and finally the delineation of compressed probabilistic state transition graphs.

The integration of all these analyses in an executable Jupyter notebook greatly eases their reproducibility, as well as the inclusion of further extensions. This notebook can further be used as a template and enriched with other ColoMoTo tools to enable comprehensive dynamical analyses of biological network models.

## 1. Introduction

Since the seminal studies from Motoyosi Sugita [1], Stuart Kauffman [2] and René Thomas [3], Boolean models have been increasingly applied to biological signalling and regulatory networks [4, 5,6, 7). With the rise of high-throughput functional genomic methods and the development of molecular pathway knowledge databases, modellers are now coping with networks encompassing hundreds of regulatory components controlling cell fate decisions for normal and pathological conditions (see e.g. [8]).

In parallel, dozens of computational tools have been developed to infer, edit and analyse the dynamical properties of Boolean networks, using different programming languages and model formats (see e.g. [9] and references therein). However, like experimental biologists, modellers are facing serious reproducibility challenges, which have been stressed in recent years (see[10], focusing on modelling studies using ordinary differential equations).

About a decade ago, several international teams working on the development of Boolean modelling tools gathered efforts to address these challenges. The aim was to delineate a common standard format to foster the exchange and reuse of qualitative models between different software tools, which took the form of a dedicated extension of a new, modular release of the popular *SBML* format, known as *sbml-qual* [11].

The second effort was put on the integration of a growing set of tools in a *Docker* container to ease their installation and articulation in flexible analysis workflows. The first release of the resulting *Common Logical Modelling Toolbox* (or *CoLoMoTo*) encompassed six complementary software tools [12]. Since then, the number of tools integrated into the CoLoMoTo suite has been constantly increasing, reaching a total of twenty in 2024. The tools currently integrated in the suite are listed in **Figure 1**, with key characteristics.

**Figure 1.**
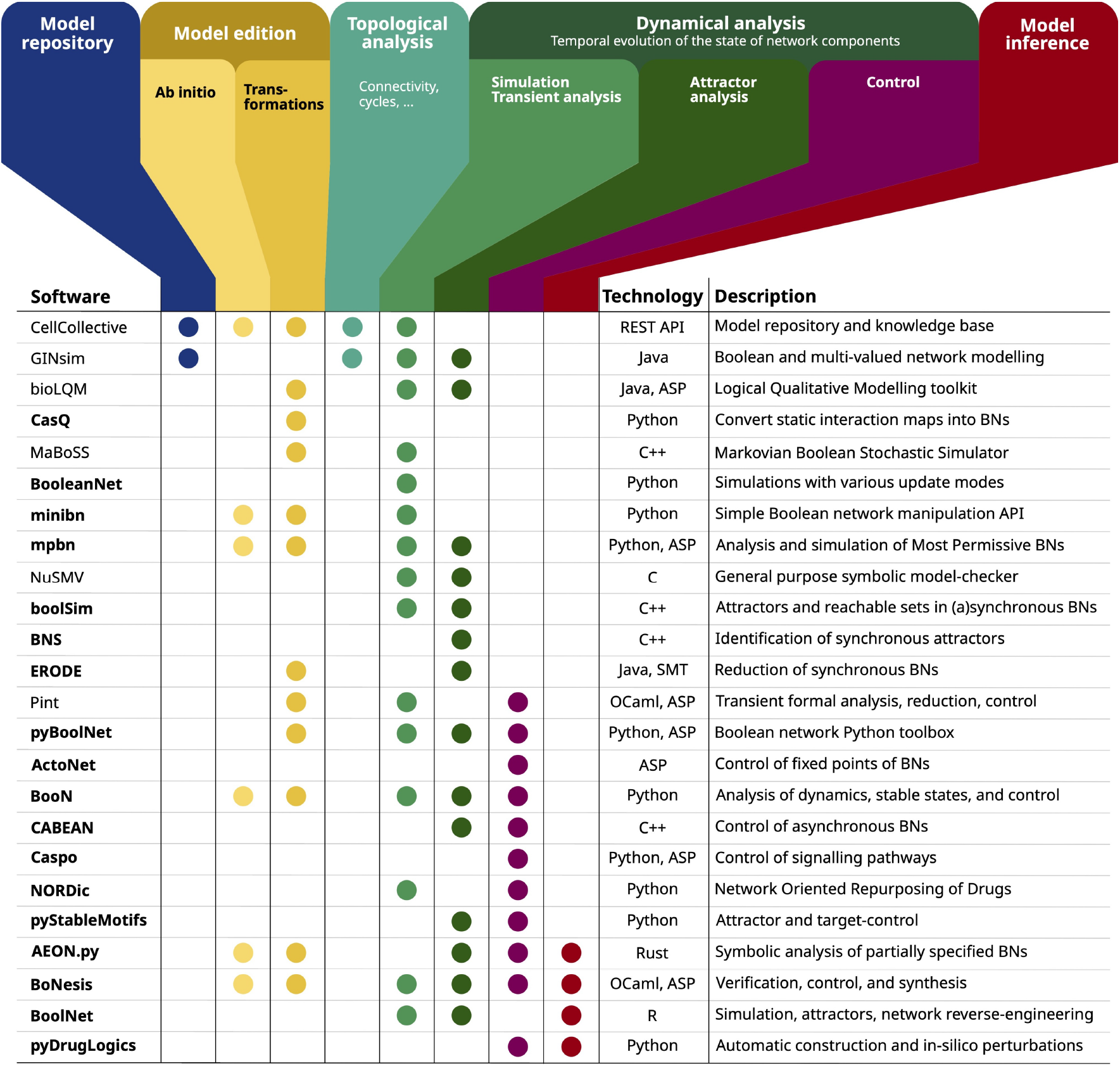
Tools integrated in the CoLoMoTo suite. Names in bold refer to tools incorporated since 2018.

In this article, we propose an up-to-date tutorial to help users take advantage of the multiple tools integrated into the *CoLoMoTo* software suite and perform sophisticated Boolean model analyses of complex signaling/regulatory networks. This tutorial relies on a comprehensive model of key signalling pathways and a core regulatory network controlling mammalian cell proliferation, published by Sizek et al. in 2019 [13].

As we shall see, the tutorial includes chunks of *python* code to reproduce several of the results and figures published in the original article and further extends these results with the help of several tools included in the *CoLoMoTo* software suite. This tutorial specifically aims to demonstrate the added value of combining different tools to perform reproducible dynamic analyses.

## 2. Material and methods

### 2.1 Installation

This notebook can be run on a recent personal computer with at least 16 Go of RAM and about 10 Go of available disk space.

The CoLoMoTo environment can be installed either as a *Docker* container or as individual *conda* packages.

#### 2.1.1. Installation of the CoLoMoTo Docker container

*Docker* and *Python* need to be pre-installed, go to your folder and then type in a terminal:

~~~
$ pip install -U colomoto-docker
$ colomoto-docker -V 2025-01-01 --bind .
~~~

Where $ stands for the terminal prompt.

#### 2.1.2. Installation of CoLoMoTo conda packages

As an alternative to using *Docker*, one can rely on *conda* to create a software environment able to reproduce the notebook on *Linux* and *macOS* computers by typing the following commands in a terminal:

~~~
$ conda create -n sizek19 -c colomoto -c potassco -c conda-forge ginsim-python bns-python boolsim-python pymaboss notebook seaborn numpy pandas matplotlib
$ conda activate sizek19
~~~

To launch this notebook, type the command:

~~~
$ jupyter notebook
~~~

Because *BNS* and *boolSim* executables are not available for *Windows®* operating systems, the complete notebook can be executed only within *Docker* on these systems.

More generally, as the versions of package dependencies cannot be fully controlled with the aforementioned *conda* command, reproducibility is better ensured by using the *Docker* container.

### 2.2. Model

The model published by Sizek et al. [13] has been imported and edited with the *GINsim* software [14]. It is publicly available in the *zginml* format (including the layout, to be open with *GINsim*, version ≥ 3.0), as well as in the sbml-qual format, in the *GINsim* model repository (http://ginsim.org/). Hence, the model can be directly loaded from the notebook using the link to the corresponding *GINsim* model repository entry.

Note that if you wish to work with this model offline, it can be downloaded locally and the loading command can be modified accordingly.

## 3. Model analysis

The following sections of the notebook cover the following analyses:

- Loading of the packages required for the tutorial
- Loading of the model and visualisation of the network with *GINsim*.
- Attractor analysis with *bioLQM*.
- Computation of synchronous attractors with *BNS*.
- Analysis of the attractors of the model modules using *bioLQM, BNS* and *boolSim*.
- Stochastic dynamical simulations of the wild type and mutant models with *MaBoSS*.
- The construction of compressed probabilistic transition graphs with *MaBoSS*.

### 3.1. Loading required packages

Before running the analyses, the packages to be used need to be loaded. For this tutorial, only the necessary CoLoMoTo packages are selected. The other packages listed in **Figure 1** can be loaded according to the instructions provided at https://colomoto.github.io/colomoto-docker/. Note that these packages can also be loaded later on, just before their use.

Pieces of code will be shown in this tutorial in boxes as follows. They represent code cells extracted from the *python jupyter* notebook.

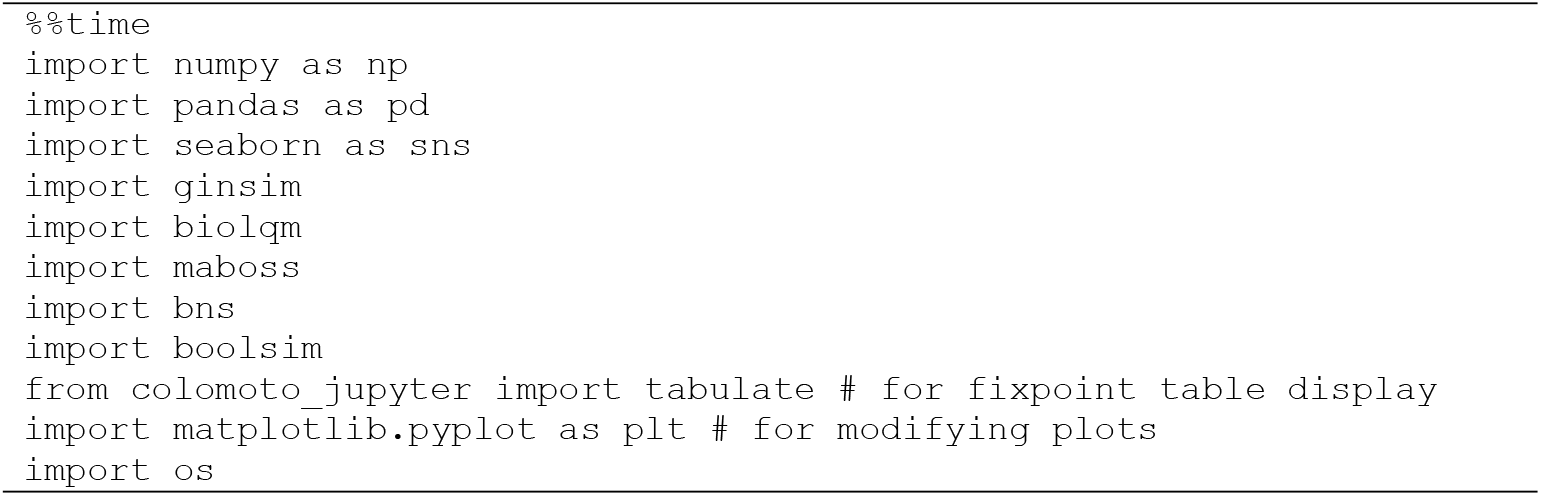

The execution of this cell outputs a few lines specifying what docker image and how much computer resources were used.

### 3.2. Loading and visualisation of the model with *GINsim*

We have used *bioLQM* [15] to import the model published by Sizek et al., and we updated the model layout with *GINsim*. The resulting model has the same logical rules as the original one and can be found in the *GINsim* repository at the address: http://ginsim.org/node/258

To load the model and visualise its *regulatory graph*, we run the following code cell:

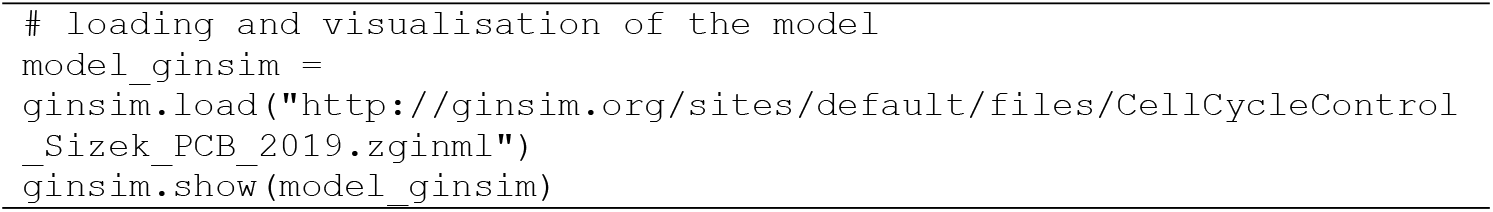

Figure 2. displays the corresponding regulatory graph.

**Figure 2.**
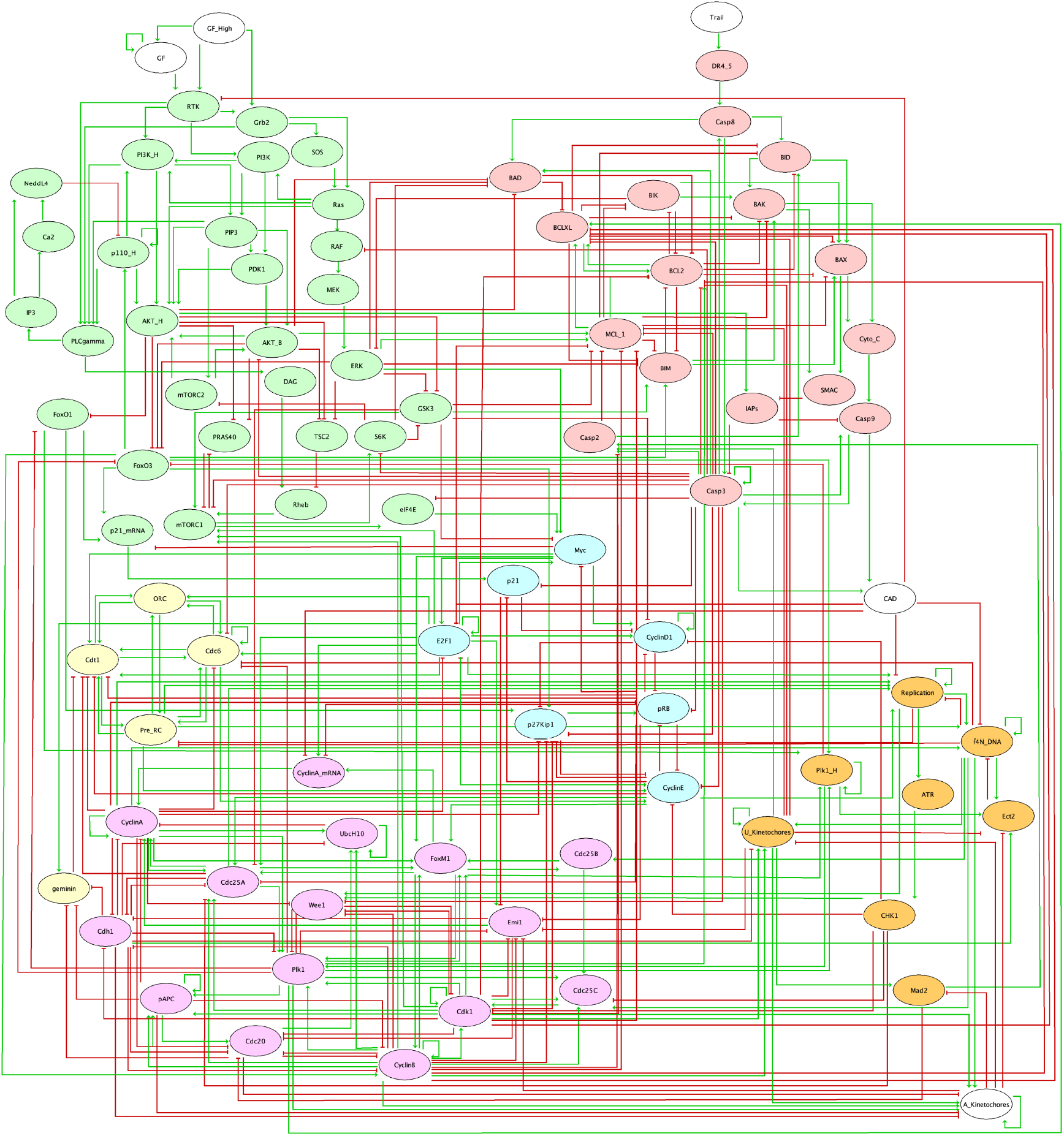
Boolean model published by Sizek et al (2019). The corresponding regulatory graph encompasses 87 components, which represent different molecular species and cellular processes involved in the regulation of cell proliferation and apoptosis in a generic human cell.

This regulatory graph encompasses 87 components representing different molecular species and cellular processes involved in the regulation of cell proliferation and apoptosis in a generic human cell. The authors conceived this complex model as an association of six main functional modules:

- The growth signaling module (green nodes) incorporates growth signaling pathways driving cell cycle commitment, responsible for modeling the dynamics of PI3K, AKT1, MAPK, and mTORC.
- The restriction switch module (blue nodes): is responsible for commitment to DNA synthesis.
- The origin licensing switch module (yellow nodes) controls licensing and firing of replication origins.
- The phase switch module (pink nodes) controls cell cycle progression (G2 -> M -> G1), taking into account the role of Polo-like kinase 1 (Plk1) during mitosis.
- The apoptosis switch module (red nodes) implements the decision between cell survival versus apoptosis.
- The cellular processes module (orange nodes) includes nodes helping to identify the phenotypic state and covering additional regulatory mechanisms.

In the regulatory graph, green arrows denote activatory interactions, while red T arcs represent inhibitions. As many model components are regulated by multiple components, their behaviour in response to regulatory input levels is encoded in Boolean rules, which combine literals (i.e., node ids) with the classical Boolean operators NOT (written “!” in GINsim), OR (“|”), and AND (“&”). The rules defined in the initial publication were encoded in the *zginml* file available in the *GINsim* model repository.

### 3.3. Attractor analysis with *bioLQM*

To obtain a first overview of the asymptotic properties of the model, we can use the software tool *bioLQM* [15], which includes an efficient algorithm to identify all stable states, regardless of their reachability [16].

The model is first converted in the *bioLQM* format, which can be achieved with the following code cell:

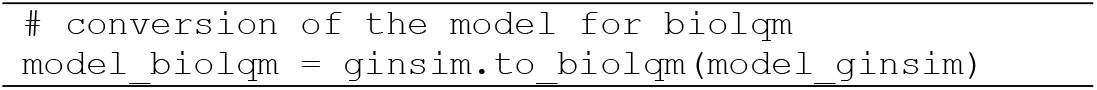

Using *bioLQM*, the *stable states* of the model can be computed in the following code cell. They correspond to long-term solutions of the model.

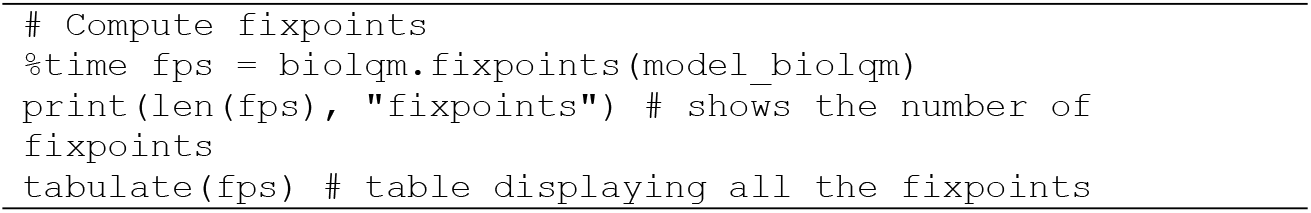

The execution of this code cell outputs a table listing the eight stable states (also called *fixed points*) of the model (shown in the notebook), matching those reported by Sizek and colleagues in Table S2 of their article.

To better visualise any of these stable states, the corresponding values can be projected on the model regulatory graph with a simple code line, using *GINsim*. Such a visualisation highlights the pathways/modules that are active in one particular stable state. An example of such stable state projection is provided in the notebook.

### 3.4. Computation of synchronous attractors with *BNS*

In their publication, Sizek et al. report the existence of a cyclic attractor when a synchronous updating method is used (i.e., when all components called to be updated by the Boolean rules at a given state are simultaneously updated, leading to a unique successor state).

*BNS* can be used to compute the synchronous attractors of the model (i.e., fixed points and simple cyclic trajectories in the synchronous case). Although *BNS* can search for attractors of any length, this search fails with such a complex model. To overcome this limitation, a range of cycle lengths is defined and attractors for the corresponding lengths are explored sequentially.

As Sizek et al. reported a synchronous cycle of length 21 in their Table S2, we make sure to include this value in the interval considered.

Of note, an attractor of length k is also an attractor of length 2k, 3k, etc. In particular, fixed points are detected as attractors of any length. Hence, in the following code cell, we consider a maximum length of 32 and filter the attractors amounting to iterations of smaller ones.

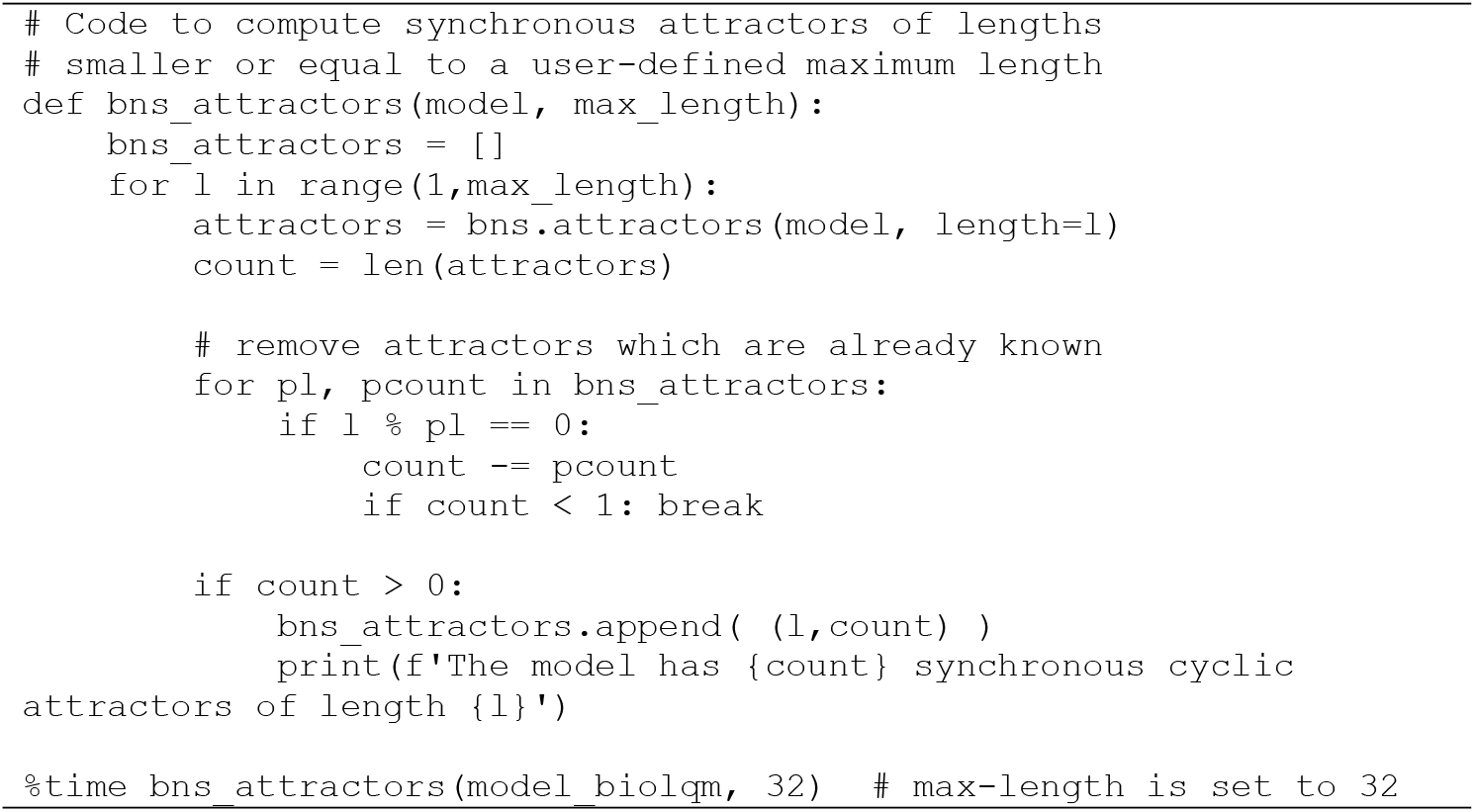

In addition to the eight fixed points already identified (corresponding to *cyclic attractors of length 1*), BNS returns four attractors of length 10, and one attractor of length 21, which correspond to the cyclic attractors reported by Sizek et al. in their supplementary Table S2.

Of note, attractors of lengths higher than 32 cannot be excluded by this analysis, as we use this number as a limit length to avoid long computations.

### 3.5. Analysis of the model constitutive modules

To generate their complex model, Sizek and colleagues initially developed smaller models for different functional modules (cf. Introduction). These modules are provided by the authors in their supplementary material and we stored this information into a zip file available on *github*. The following code cell retrieve this information and computes the total number of components, the number of other components regulating the components of the module (aka *inputs*), the number of stable states, the number of terminal trap spaces (comprising the stable states, but potentially also approximations to cyclic attractors), as well as the numbers of synchronous and asynchronous attractors for each of these modules.

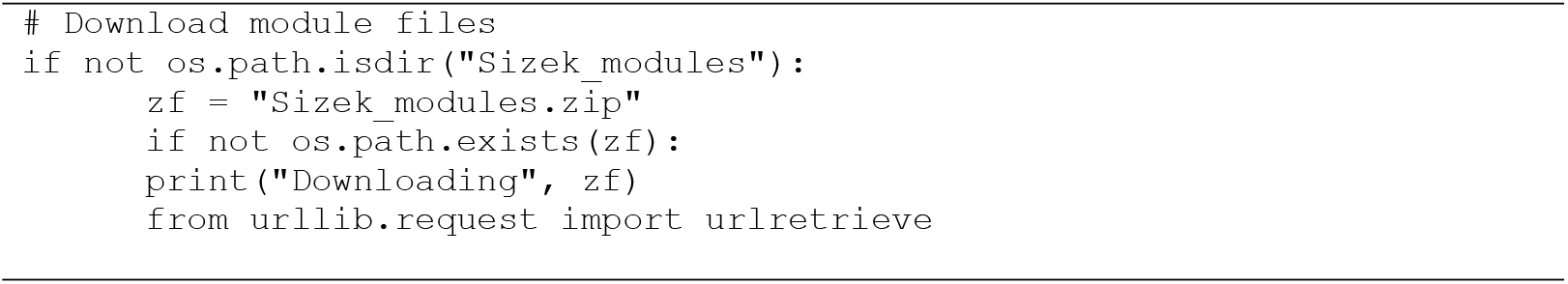
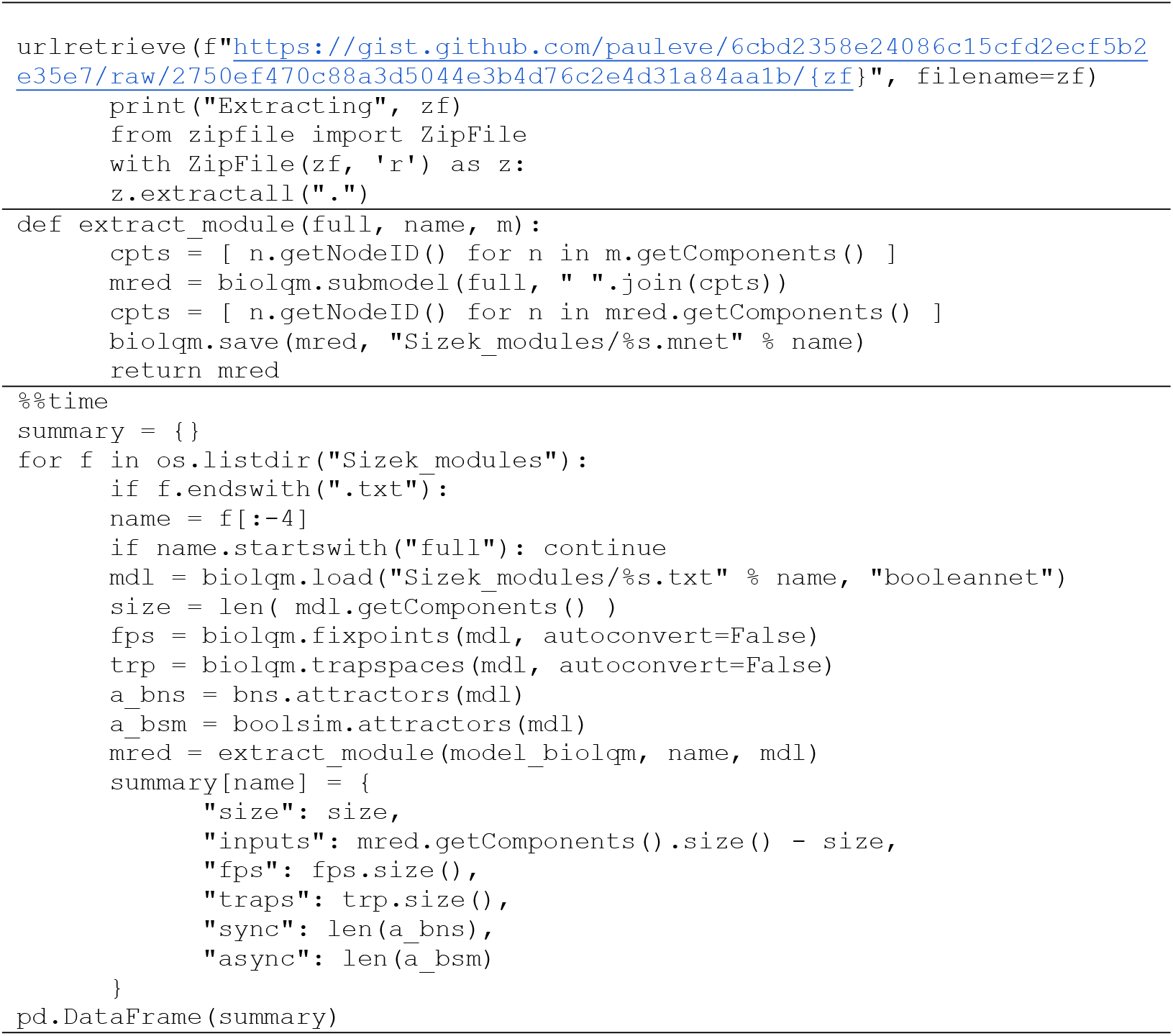

The execution of this code cell generates the results reported in **Table 1**.

**Table 1.**
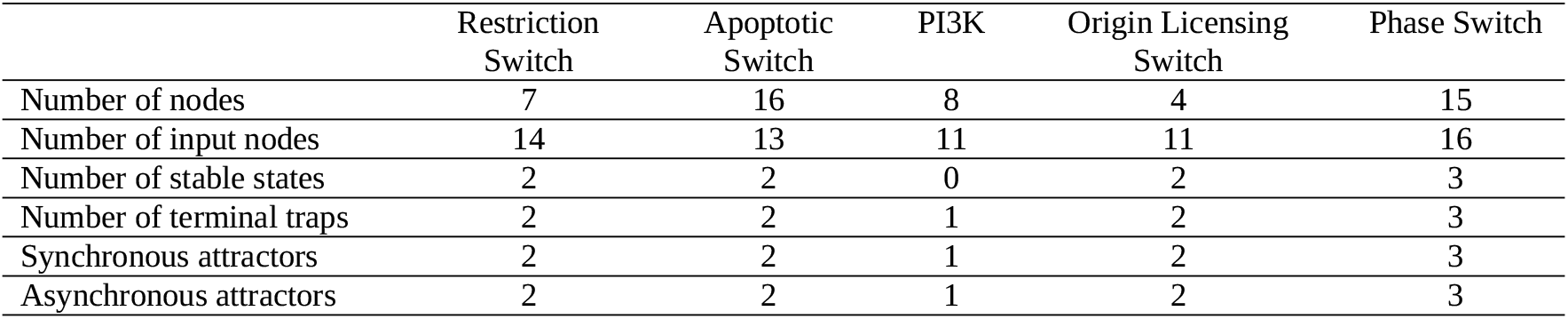
Some properties of the modules constitutive of the model of Sizek et al. (2019)

|  | Restriction<br>Switch | Apoptotic<br>Switch | PI3K | Origin Licensing<br>Switch | Phase Switch |
| --- | --- | --- | --- | --- | --- |
| Number of nodes | 7 | 16 | 8 | 4 | 15 |
| Number of input nodes | 14 | 13 | 11 | 11 | 16 |
| Number of stable states | 2 | 2 | 0 | 2 | 3 |
| Number of terminal traps | 2 | 2 | 1 | 2 | 3 |
| Synchronous attractors | 2 | 2 | 1 | 2 | 3 |
| Asynchronous attractors | 2 | 2 | 1 | 2 | 3 |

We can observe that the modules are highly interconnected, as they all have high numbers of inputs relative to their sizes, due to regulations exerted by nodes belonging to other modules.

Furthermore, all these modules but one display multistability but no cyclic behaviour. The PI3K module is the exception, as it has a unique, cyclic attractor, regardless of the use of synchronous or asynchronous updating. The cyclic behaviour of the PI3K module is indeed reported and discussed by Sizek et al.

At this point, it becomes apparent that the global cyclic behaviour corresponding to the cell cycle (see section 8) arises from the interconnections of multiple modules.

### 3.6. Characterisation of synchronous cyclic properties with *bioLQM*

To visualise the 21 states composing the synchronous cyclic attractor, *bioLQM* can be used to generate a trace from one of the cycle states reported by Sizek et al., considering a number of steps somewhat greater than 21. First, we define an initial state belonging to the cyclic attractor, according to Sizek et al. with the following code cell:

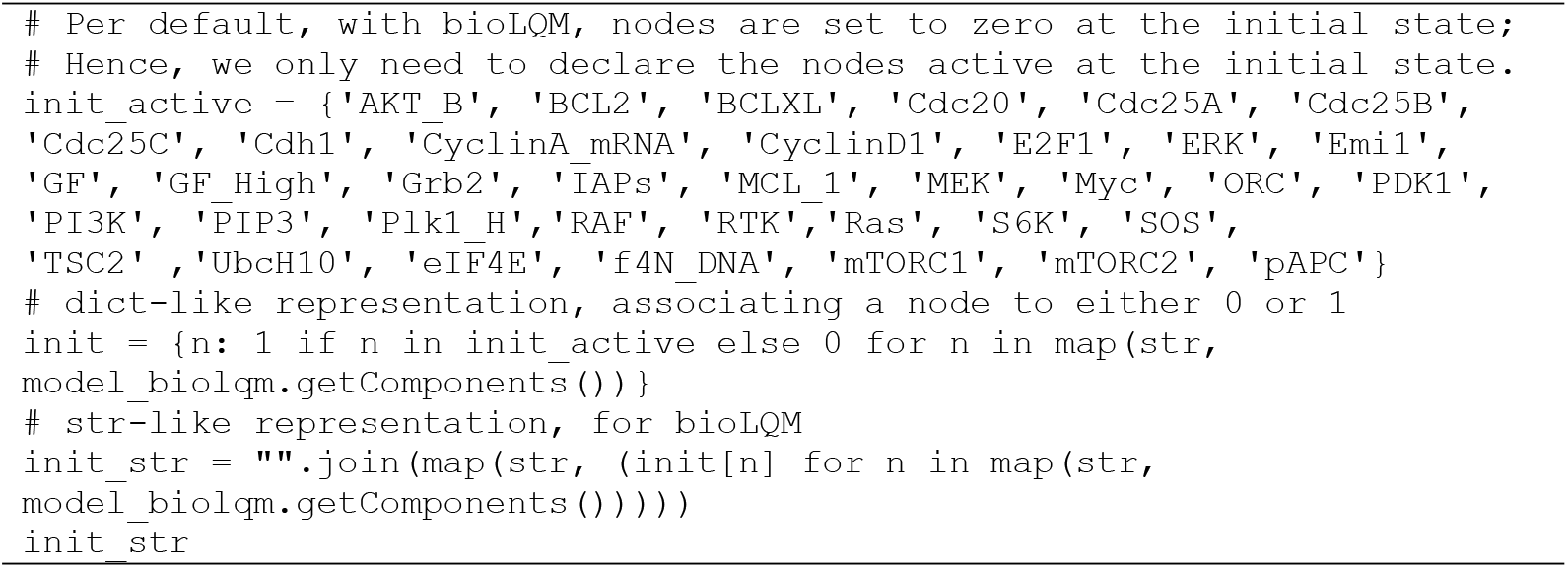

Next, we compute a synchronous updating trace, starting with this initial state and considering a maximum of 50 steps, with the following code cell:

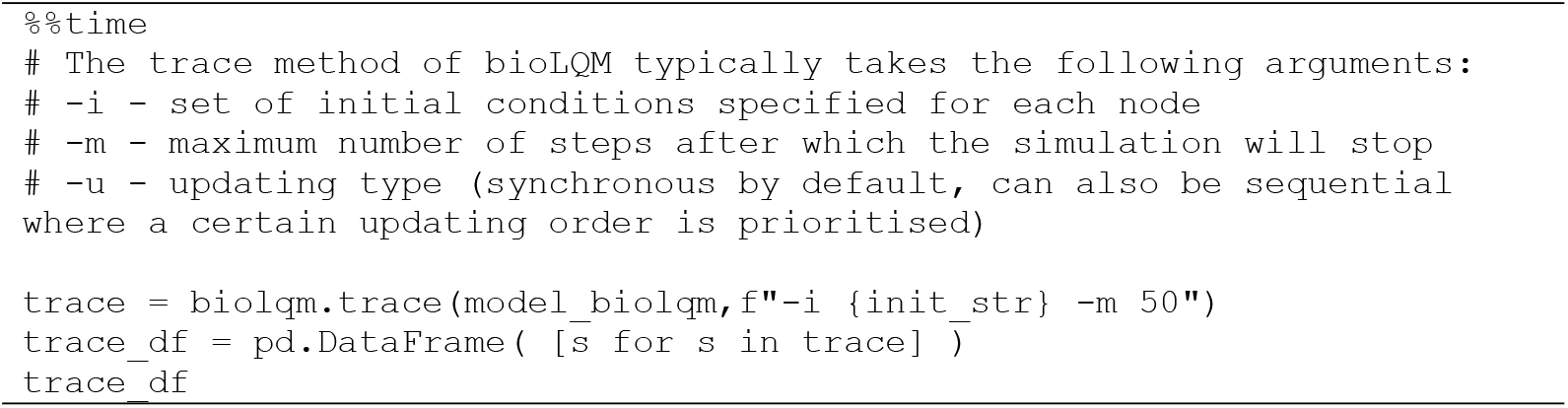

The execution of this cell generates a table encompassing 22 rows (cf. notebook), each corresponding to one state belonging to a cyclic sequence. We can easily check that the first and last states listed in the table above are indeed identical with the following code cell:

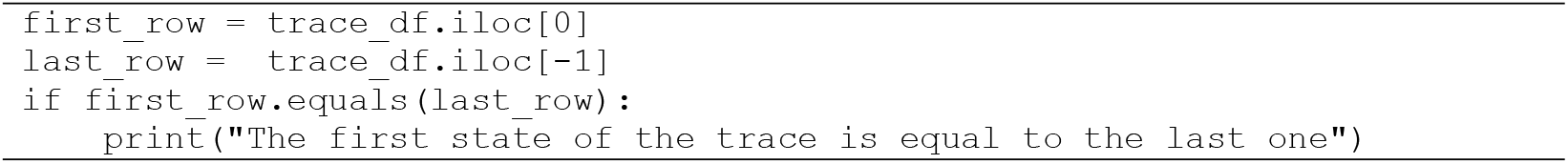

The execution of this cell confirms that we have identified the cyclic attractor of length 21 mentioned above. To ease the comparison of this sequence of states with the periodic pattern reported by Sizek et al. in their Figure 4, we can plot it with similar graphical conventions, using the following code cell:

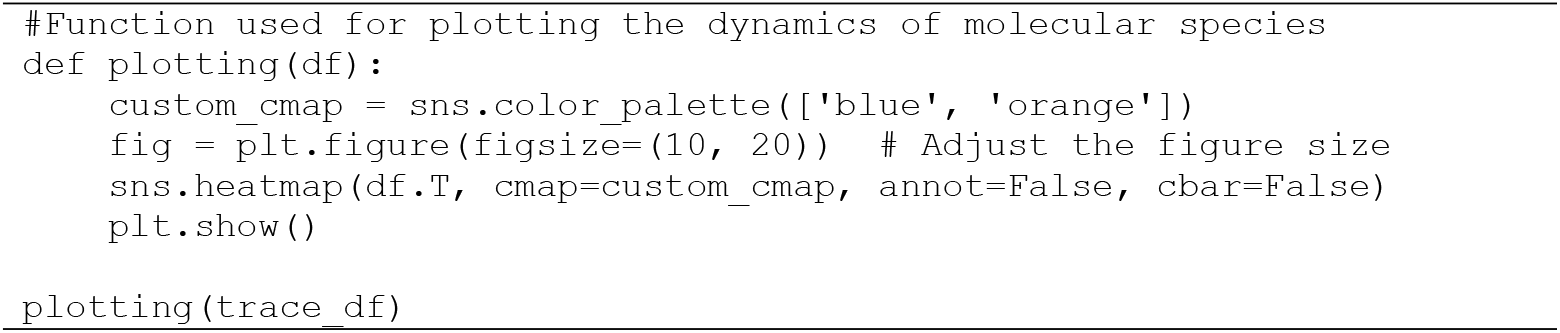

The resulting heatmap is shown in **Figure 3**.

**Figure 3.**
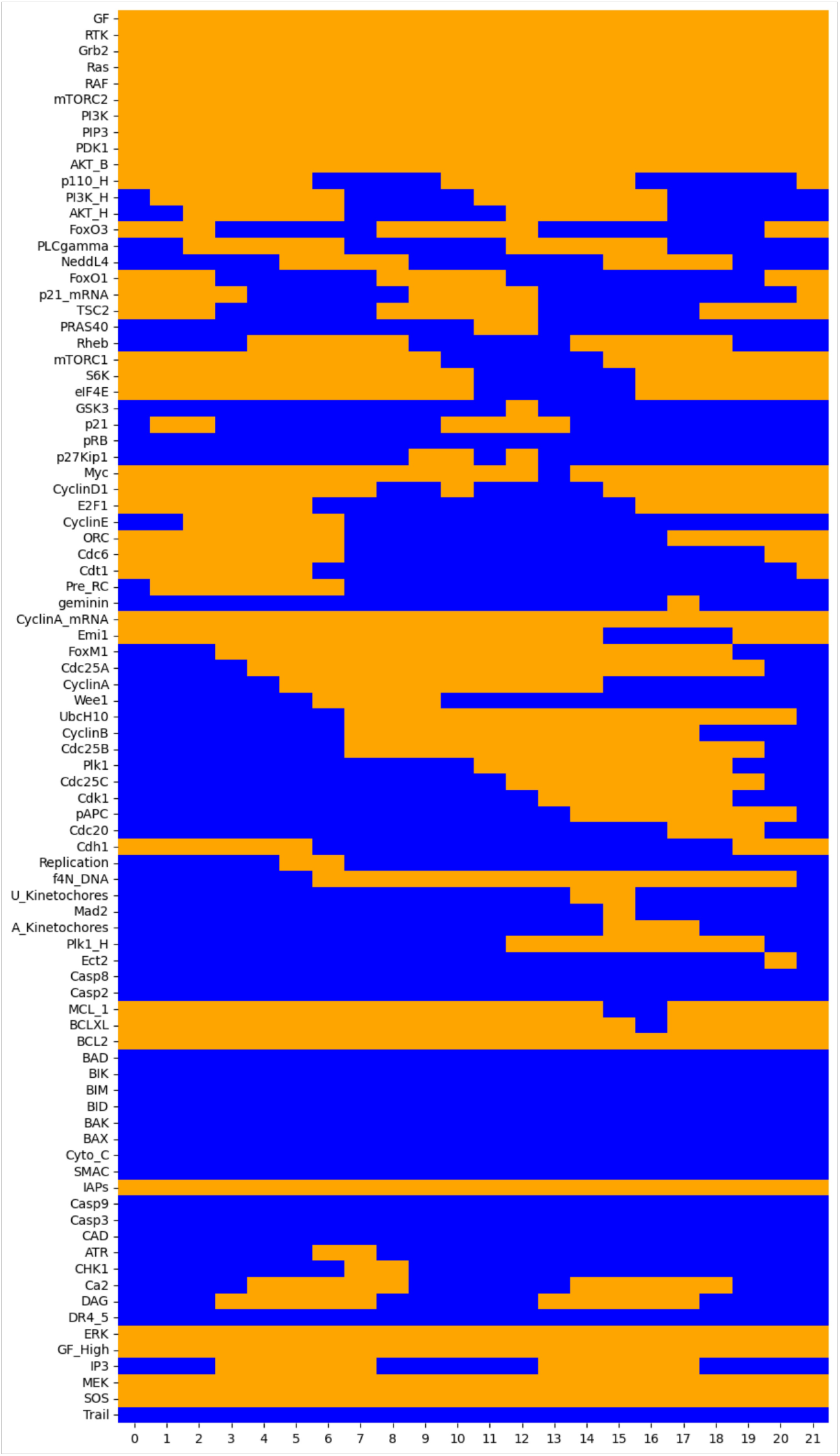
Heat map displaying the sequence of 22 states (columns) forming the synchronous cycle of length 21, with the initial state (first column, index 0) matching the last one (column with index 21). Orange cells denote the active components (rows) at the corresponding cycle state (column), whereas the blue cells denote the suppressed components.

This heat map displays a sequence of 22 states (columns) corresponding to the synchronous cycle of length 21, with the initial state (first column, index 0) matching the last one (column index 21). Orange cells denote active components (rows) at the corresponding cycle state (column), whereas blue cells denote suppressed components. Note that some nodes are never activated along the cycle, while others are always activated. In particular, the apoptotic pathway (BIM, BAX, BAK, CytC, Casp3, Casp9, etc.) is inactive in this sequence, while the cell cycle is activated by the MAPK pathway (RTK, Ras, Raf, PI3K, etc.).

Hence, using different tools, we could also identify the synchronous attractor reported by Sizek et al., which recapitulates the typical sequence of (in)activation of cell components, including cyclins. Further analysis of the timing of cyclins (in)activations is provided in section 11 below.

Of note, Sizek et al. reported that this cyclic behaviour is sensitive to the updating strategy used. To further characterise this sensitivity, *bioLQM* can be used to compute traces starting from the same initial state but using random asynchronous updating. This can be achieved using the following code cell:

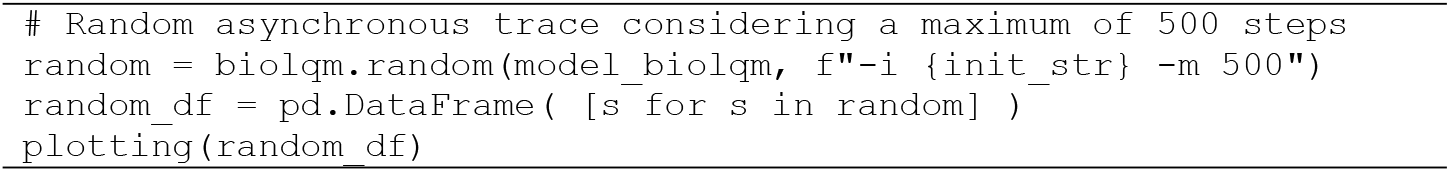

Note that a longer maximal number of steps (500) is considered here because we do not know if the trace will end up in an asynchronous cycle, or how long such a cycle might be. The resulting heatmap is shown in **Figure 4**.

**Figure 4.**
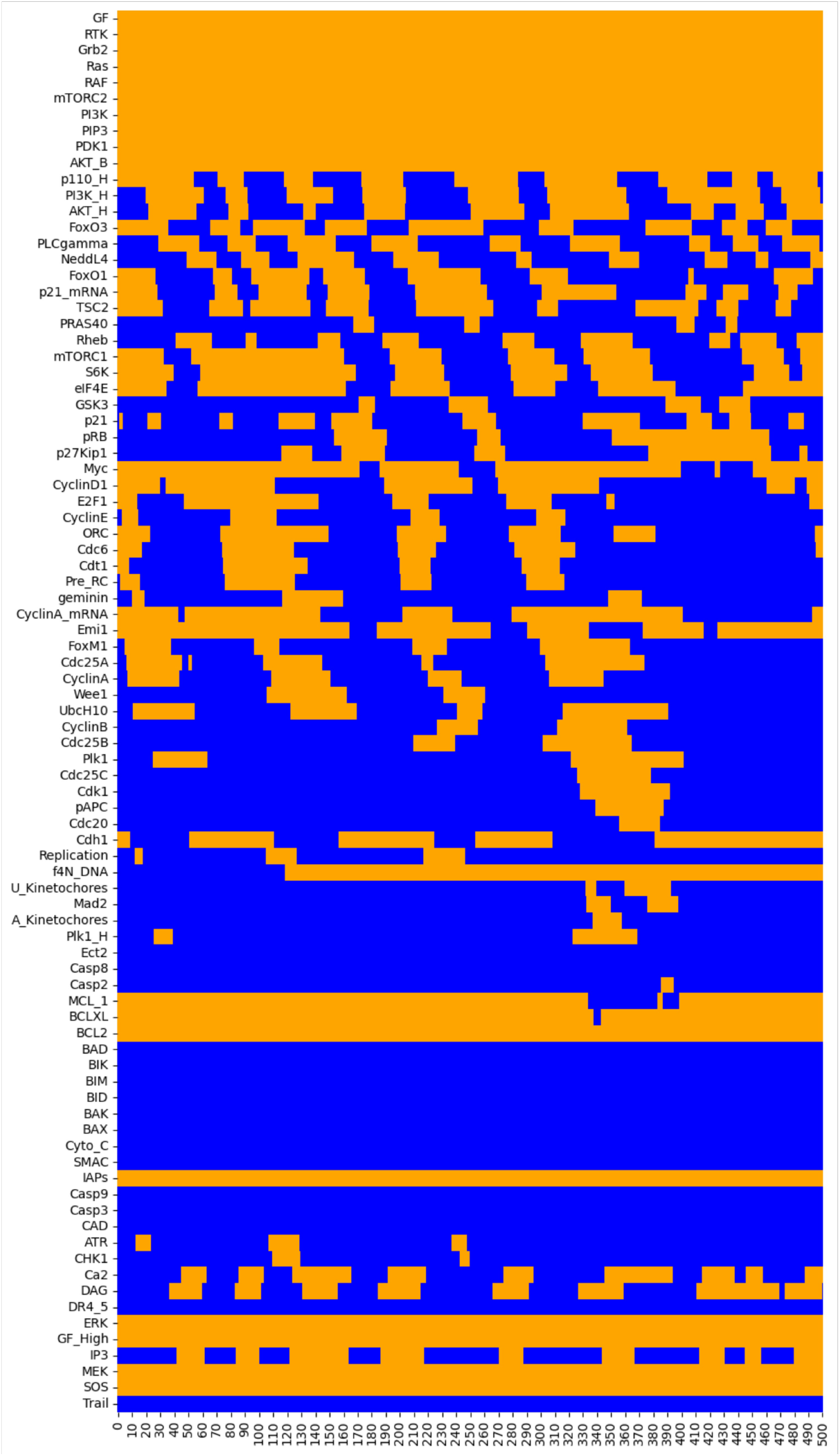
Heat map displaying a sequence of 500 states (columns) resulting from a random asynchronous updating of the model starting with the same initial state shown in Figure 3. Orange cells denote the active components (rows) at the corresponding cycle state (column), whereas the blue cells denote the suppressed components.

In this heatmap, we can observe a quasi-periodic pattern, but this pattern gets progressively degraded along the simulation. This is due to the random choice of transitions whenever several components are called to switch their values at a given state, according to their logical rules. Of note, if we repeat the same random asynchronous simulation, we typically end up with a different quasi-periodic pattern.

Interestingly, when bigger enough maximal step numbers are considered, simulations ultimately lead to a stable state corresponding to an apoptotic fate. To visualise this, in the notebook, we provide the code for a simulation using a maximum number of steps set to 2000, which needs to be uncommented to be run.

In their manuscript, the authors defined a biased transition order to enforce a more robust cyclic behaviour. Hereafter, we propose an alternative approach to deepen our understanding of the asynchronous behaviour of the model, taking advantage of the software *MaBoSS* [17].

### 3.7. Using *MaBoSS* software to perform stochastic simulations of the wild type model

*MaBoSS* performs stochastic simulations over Boolean networks based on the continuous-time *Markov chain* formalism, relying on the *Gillespie algorithm* [17]. The software enables the generation of *time plots* showing the evolution of mean component values (or patterns thereof) and *pie charts* showing the probabilities of the different model states reached at the end of the simulation.

The *bioLQM* model can be converted into the format required by *MaBoSS* using *bioLQM* with the following code cell:

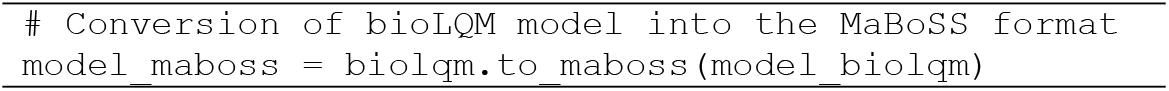

After loading the model, simulation parameters can be modified with the following code cell:

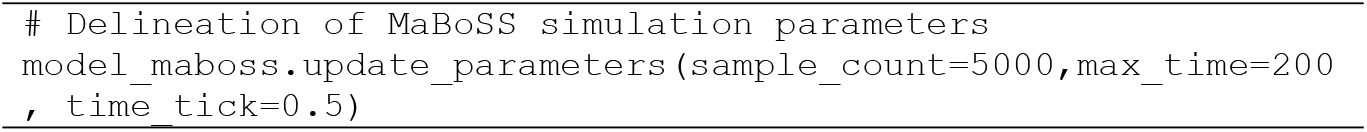

This code cell defines the following simulation parameters:

- Number of simulations: 5000.
- Maximum length of the simulation (in arbitrary unit): 200.
- Interval between two time points: 0.5.

These simulation parameters will be used for all the stochastic simulations reported in the notebook.

Next, we define a wild-type version of the model, called *WT*, and we select the *external variables*, i.e., the variables for which the values will be reported during the simulation. This is crucial to avoid combinatorial explosion.

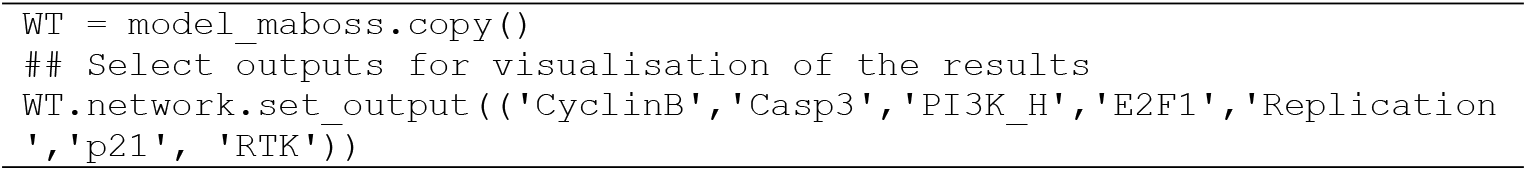

Note that when exporting a model to *MaBoSS* with *bioLQM* or *GINsim*, all the initial values are set to 0 by default. Hereafter, we assign to the model the initial conditions defined by Sizek et al. in their Figure 5.

**Figure 5.**
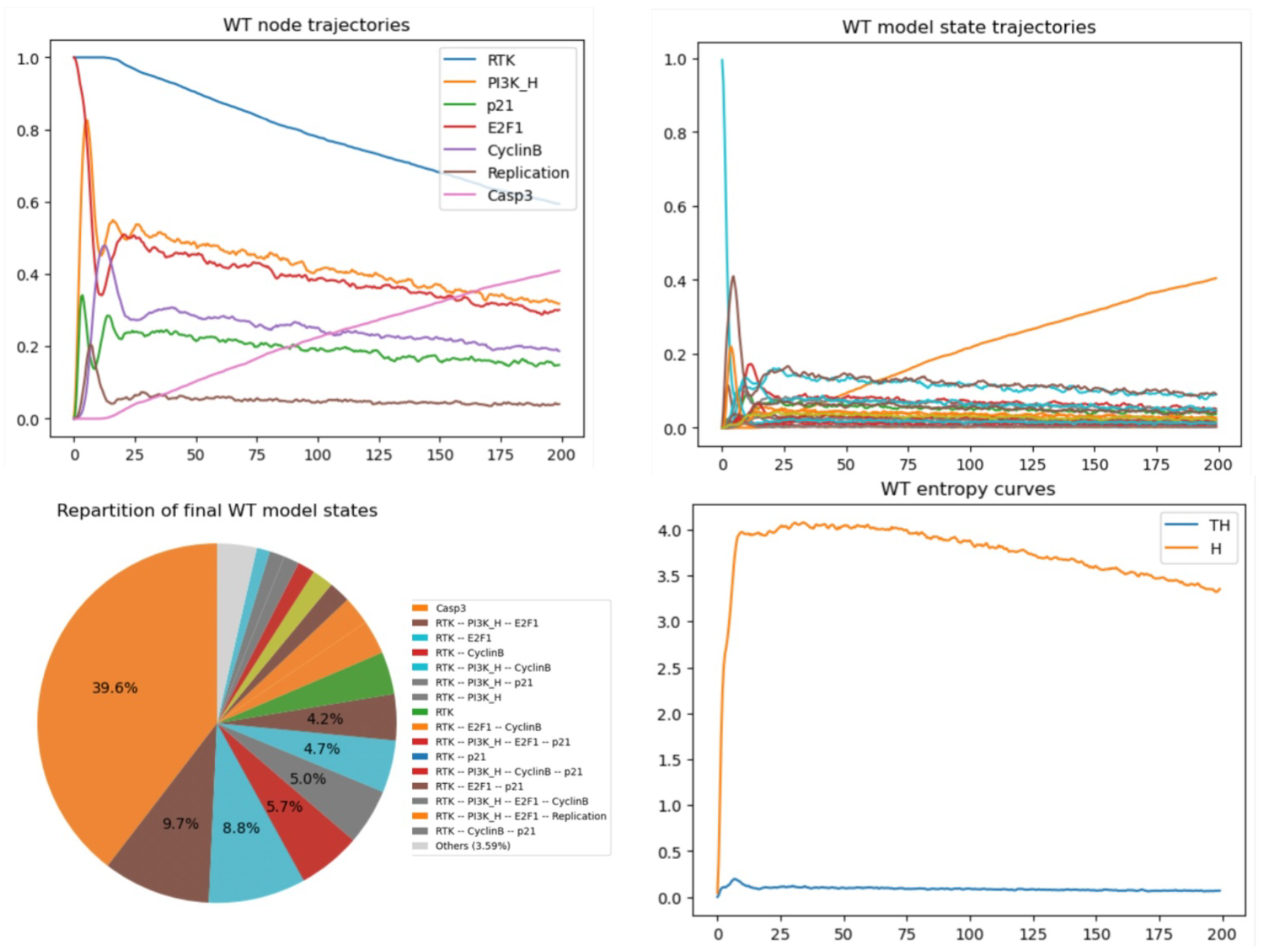
*MaBoSS* simulations of the wild-type model. (Upper left panel) Trajectories for a subset of nodes. (Upper right panel) Trajectories for a subset of model states (legend in the lower left panel). (Lower left panel) Pie chart representing the probabilities for the model states at max time (time = 200). (Lower right panel) Entropy and transition entropy time plots.

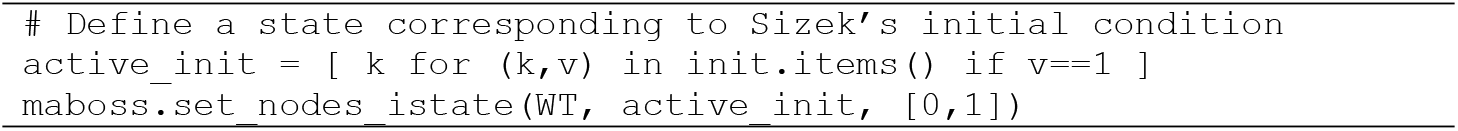

*MaBoSS* simulation results can be graphically displayed as *time plots* showing the probabilities of active nodes or model states (vectors of active nodes) over time, or as *pie charts* showing the probabilities of the final states (i.e., the states reached at the end of the simulations).

It is also possible to compute the *state entropy* (H) and the *transition entropy* (TH) over time, which can be interpreted as signatures for the stable states or cyclic attractors.

The following cell code launches a simulation of the WT model with *MaBoSS*, using the parameters defined above:

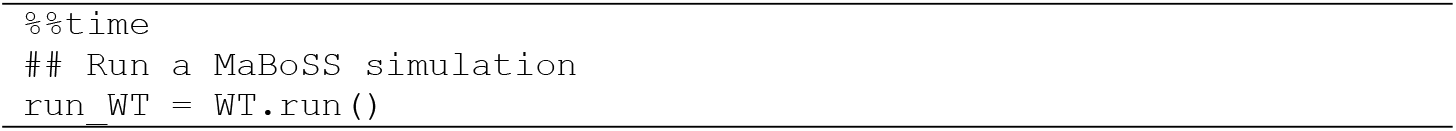

To graphically visualise the results of this simulation in terms of node or model states trajectories, of final model state repartition, and of entropy curves, we can use the following code cell:

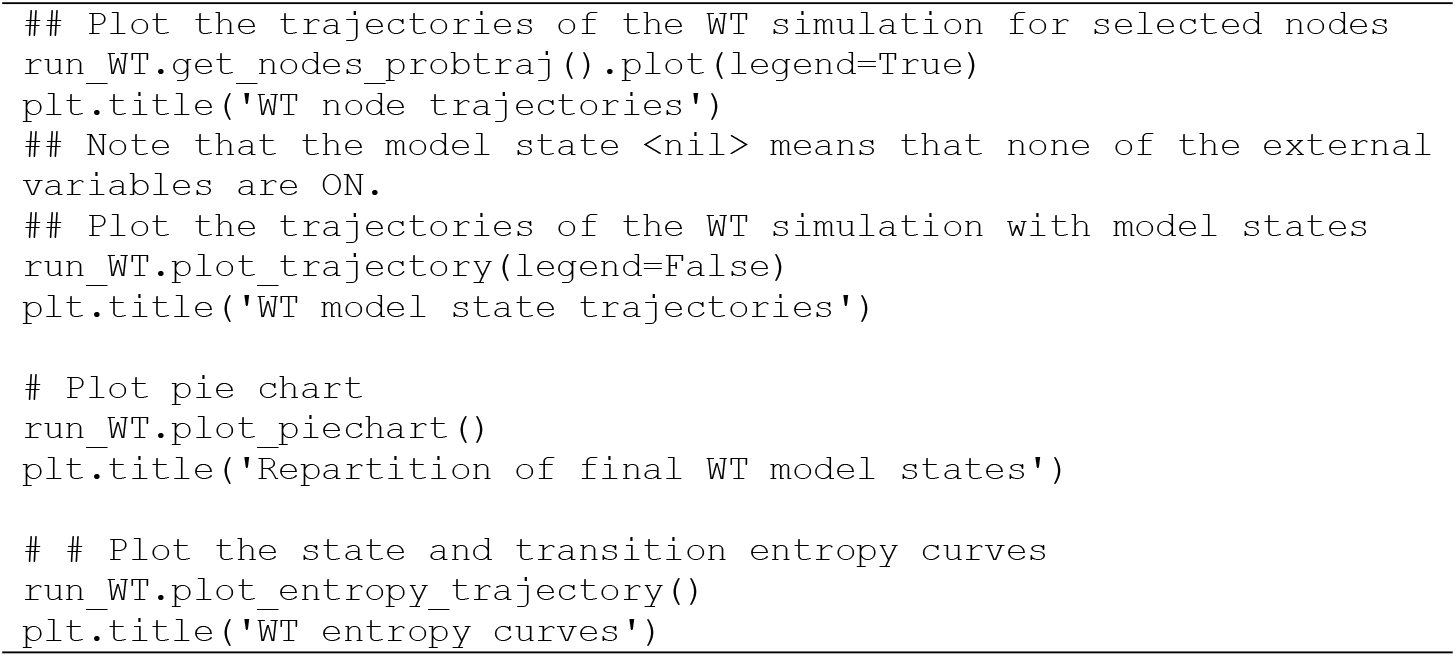

The resulting graphs are shown in **Figure 5**.

The simulations reveal a transient cyclic-like behaviour, which is eventually lost due to the spontaneous Casp3 activation, thereby revealing the progressive dominance of the apoptotic stable state.

The state entropy initially rises abruptly, and later slowly decreases. In parallel, the transition entropy curve displays a little peak before remaining close to zero, with some noise. Together, these two curves are suggestive of a quasi-periodic behaviour, which should collapse in the long term, as more and more cells trigger apoptosis.

Next, we can use *MaBoSS* to study the impact of changes in the initial conditions on the dynamic behaviour of the model. In the preceding simulation, all death signals were initially OFF. For example, we can assess the effect of activating TRAIL, keeping the rest of the other initial conditions unchanged. The following code cell defines a copy of the WT model, triggers TRAIL level to 1 at the initial state, performs the simulation, and finally displays the results:

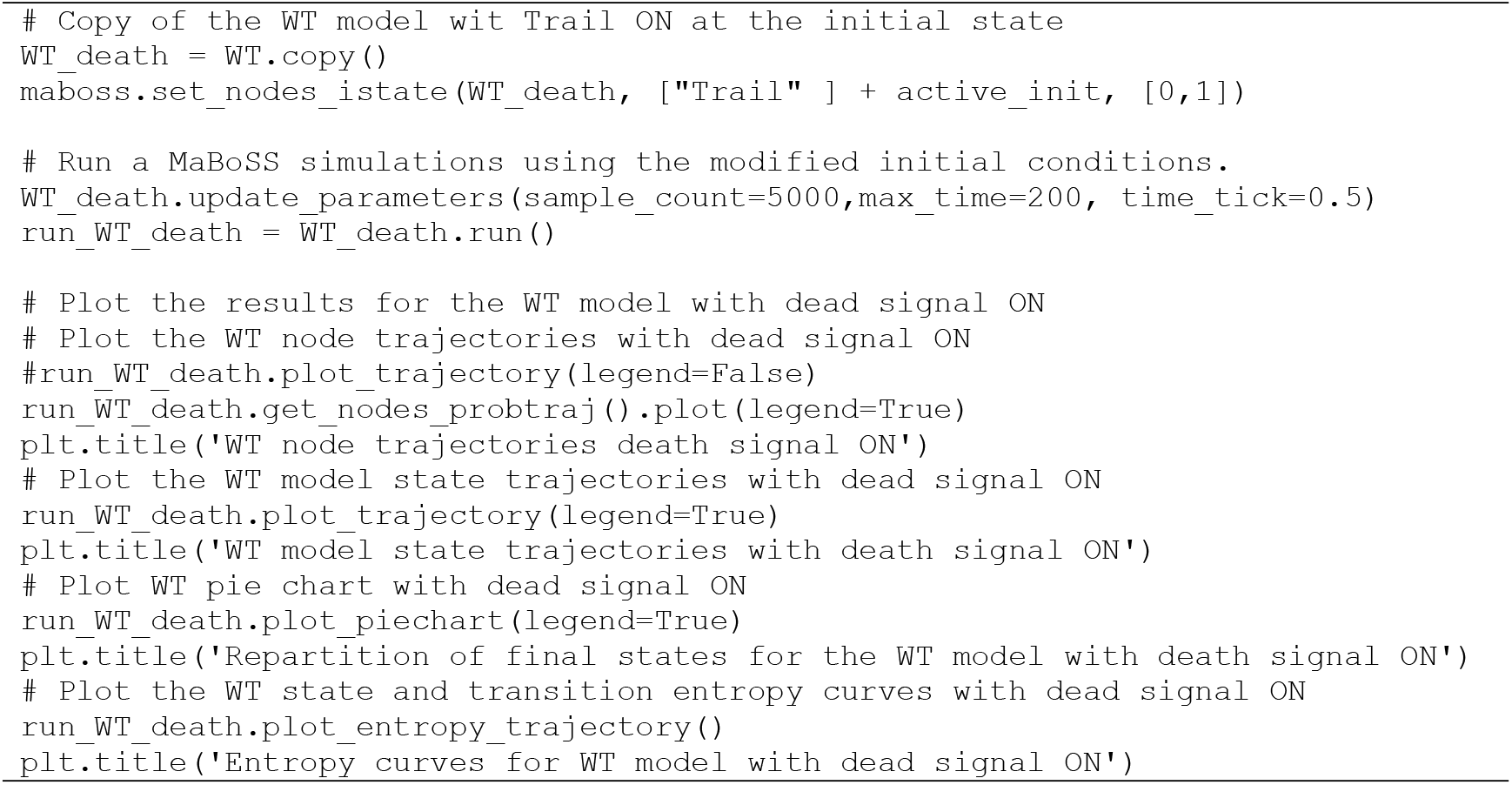

The resulting graphs are displayed in **Figure 6**.

**Figure 6.**
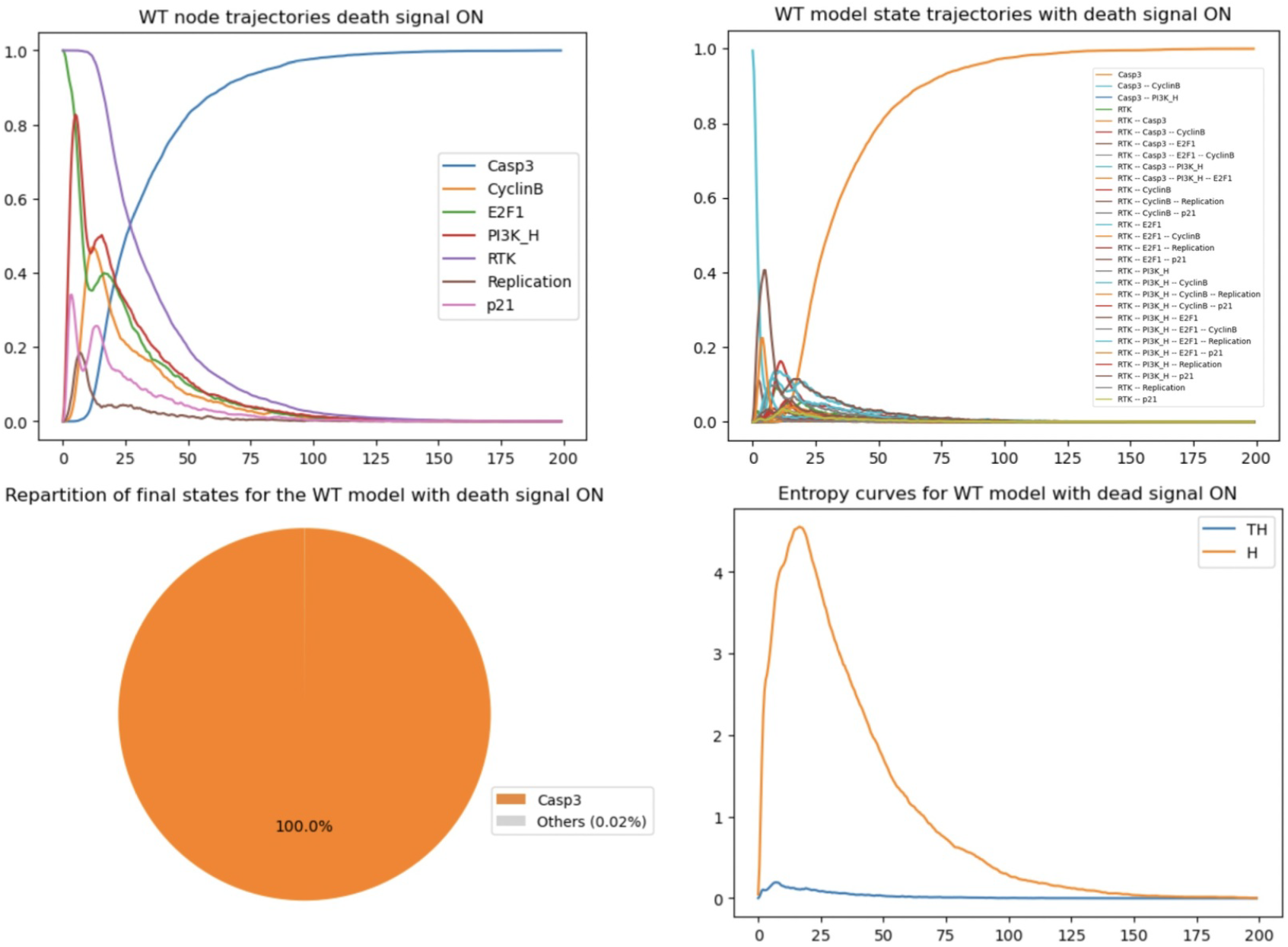
MaBoSS simulations of the wild-type condition with death receptor signals active. (Upper left panel) Trajectories for a subset of nodes. (Upper right panel) Trajectories for a subset of model states (legend in the lower left panel). (Lower left panel) Pie chart representing the probabilities for the model states at max time (time = 200). (Lower right panel) Entropy and transition entropy time plots.

Looking at the graphs of Figure 6, we can see that the activation of Trail at the initial state enables a faster rise of Casp3 and thus of the apoptotic fate. In parallel, the state entropy drops much faster, while the transition entropy stabilises around zero, suggesting a stable state situation.

### 3.8. Mutant simulations with *MaBoSS*

In their article, Sizek and colleagues report a sophisticated analysis of the role and dynamical properties of the PI3K/AKT1 pathway, and recapitulate deleterious effects of its alterations. They further analyse the impact of the blocking of the Polo-like kinase 1 (Plk1, a mitotic driver and chemotherapy target) on cell cycle progression.

Hereafter, we focus on the analysis of the impact of the perturbation of other model components, using the *MaBoSS* software.

#### 3.8.1. Stochastic simulation of Casp8 ectopic activity

Using the following code cell, we first simulate the impact of a Casp8 ectopic activity and display the resulting time plot and pie chart.

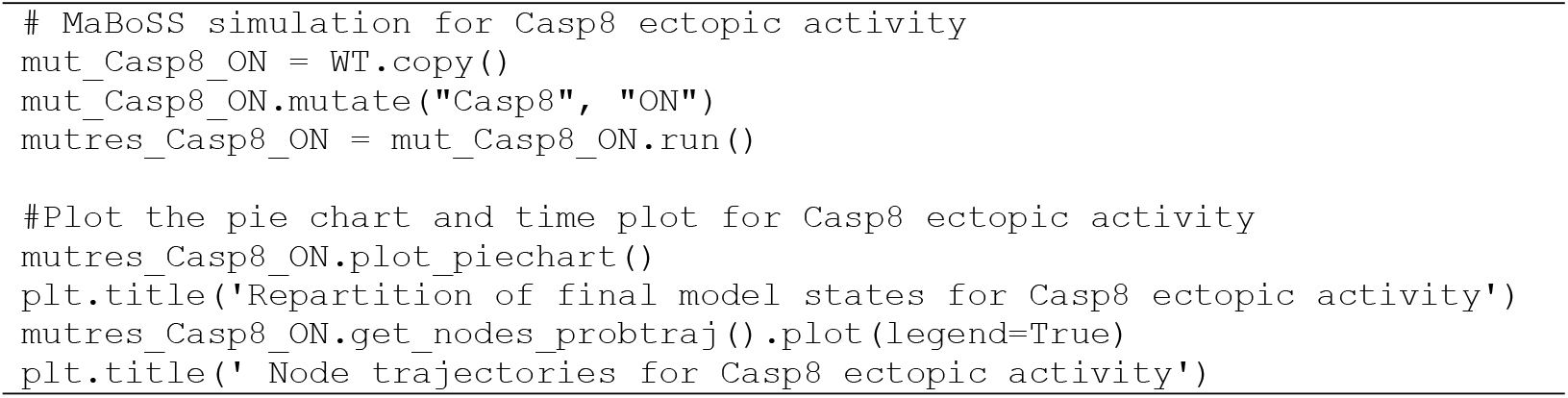

**Figure 7 (left)** shows the resulting plots. Based on these results, we can conclude that the ectopic expression of Casp8 drives the cells into apoptosis faster, no matter if TRAIL is ON or OFF.

**Figure 7.**
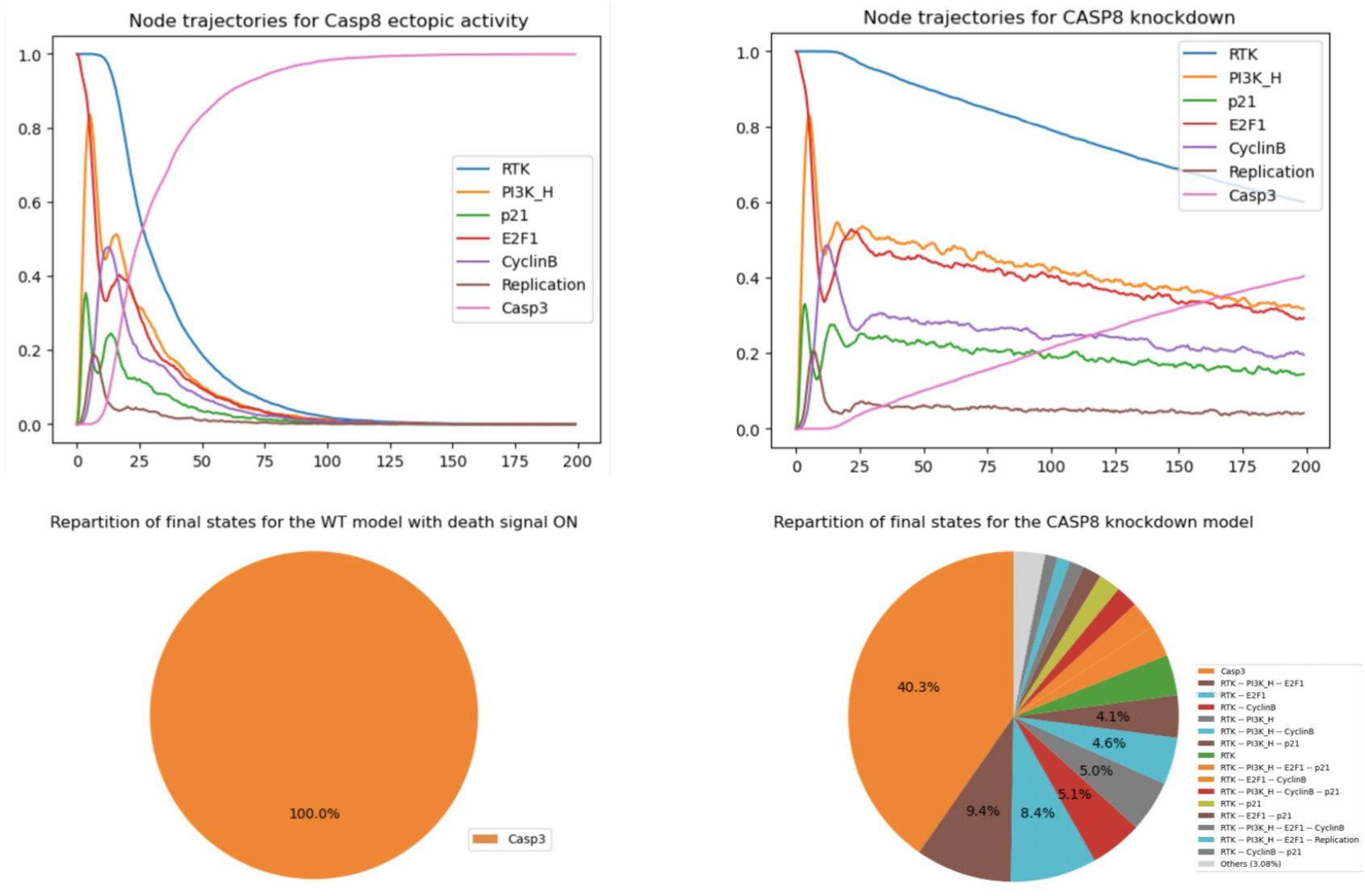
*MaBoSS* simulations for Casp 8 mutants. (Left panels) Node trajectories and pie chart showing the asymptotic solutions for Casp8 overexpression. (Right panels) Node trajectories and Pie chart representing the probabilities for the model states at the end of simulations for Casp8 knock down.

#### 3.8.2. Stochastic simulation of Casp8 knockdown

We can also check the impact of a full knockdown of Casp8 by executing the following code cell:

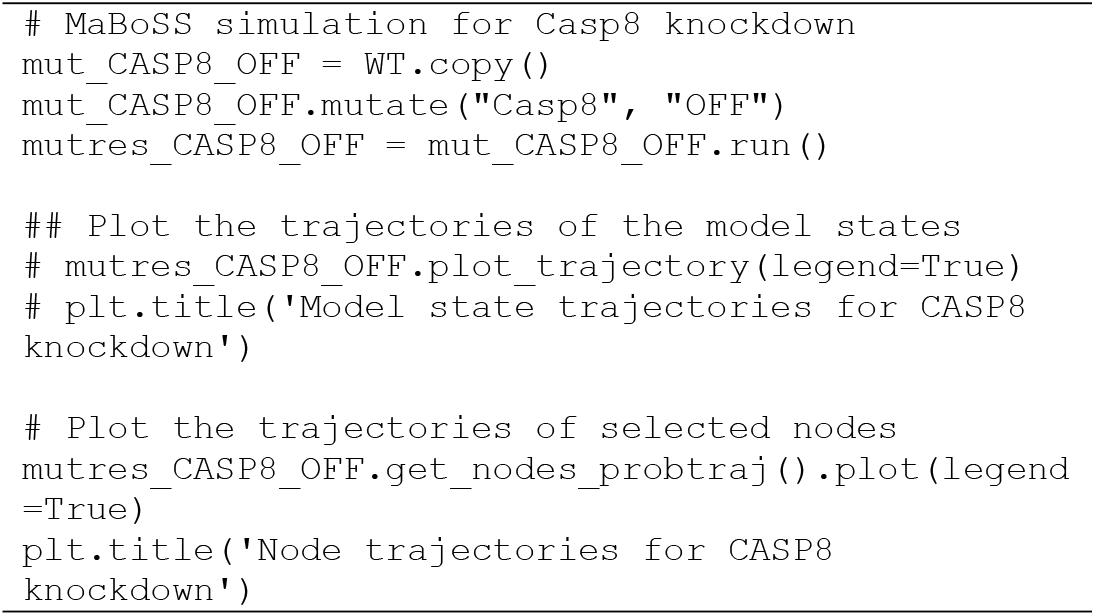
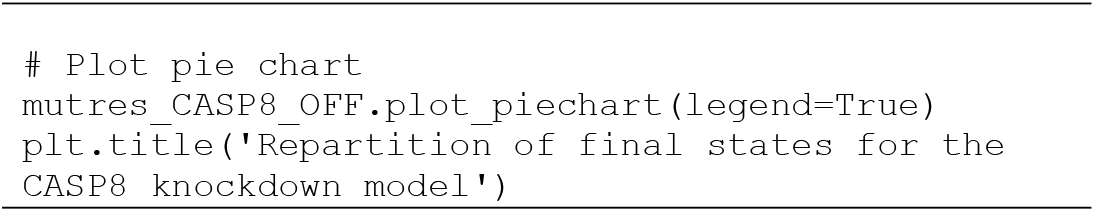

As shown in **Figure 7 (right)**, Casp3 is initially blocked but becomes ultimately activated, like in the wild-type case, independently of the status of Casp8. Hence, the model predicts that apoptosis should ultimately occur even in the absence of Casp8 activity. The knockout of Casp8 does not affect the model in the absence or presence of TRAIL.

Of note, it is possible to knock out a specific interaction with *bioLQM*, using the syntax: “target:regulator%0”, denoting a blockade of the interaction exerted by the regulator onto the target (an example is provided in the notebook).

Using a bit of *python* code, it is further possible to systematically explore a series of mutants, and to filter the results. In this respect, the notebook provides an example of such code, considering ten mutant simulations (inhibition and activation of five different nodes), and filtering those enabling a blockade of apoptosis, denoted by the suppression of Casp3 activation, while keeping the cell cycle active.

### 3.9. Deeper analysis of the quasi-cyclic synchronous attractor

In this section, we explore the dynamics of the cell cycle by studying the succession of cyclin activations and inactivations. In the WT situation, we expect that the cyclins get activated in the following order: CyclinE, followed by CyclinA, then CyclinB, and that they get inactivated as follows: Cyclin E is inactivated, then CyclinA, and finally CyclinB.

#### 3.9.1. Verification of the order of activations of cyclins in the WT situation by plotting a compressed state transition graph

The following code cell defines a simulation of the WT model focusing on cyclin dynamics.

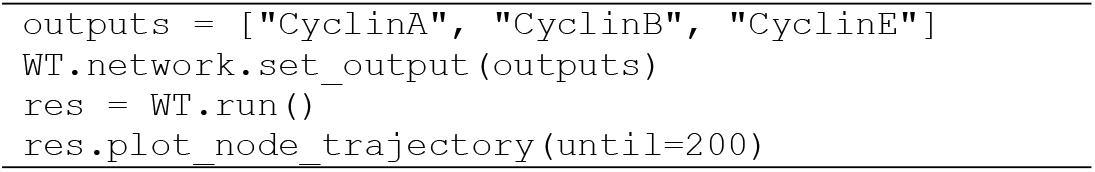

The resulting time plot is shown in **Figure 8 (top left)**. We can see that the cyclins follow the expected order. Note that as *MaBoSS* reports mean probabilities over time, these oscillations are rapidly damped.

**Figure 8.**
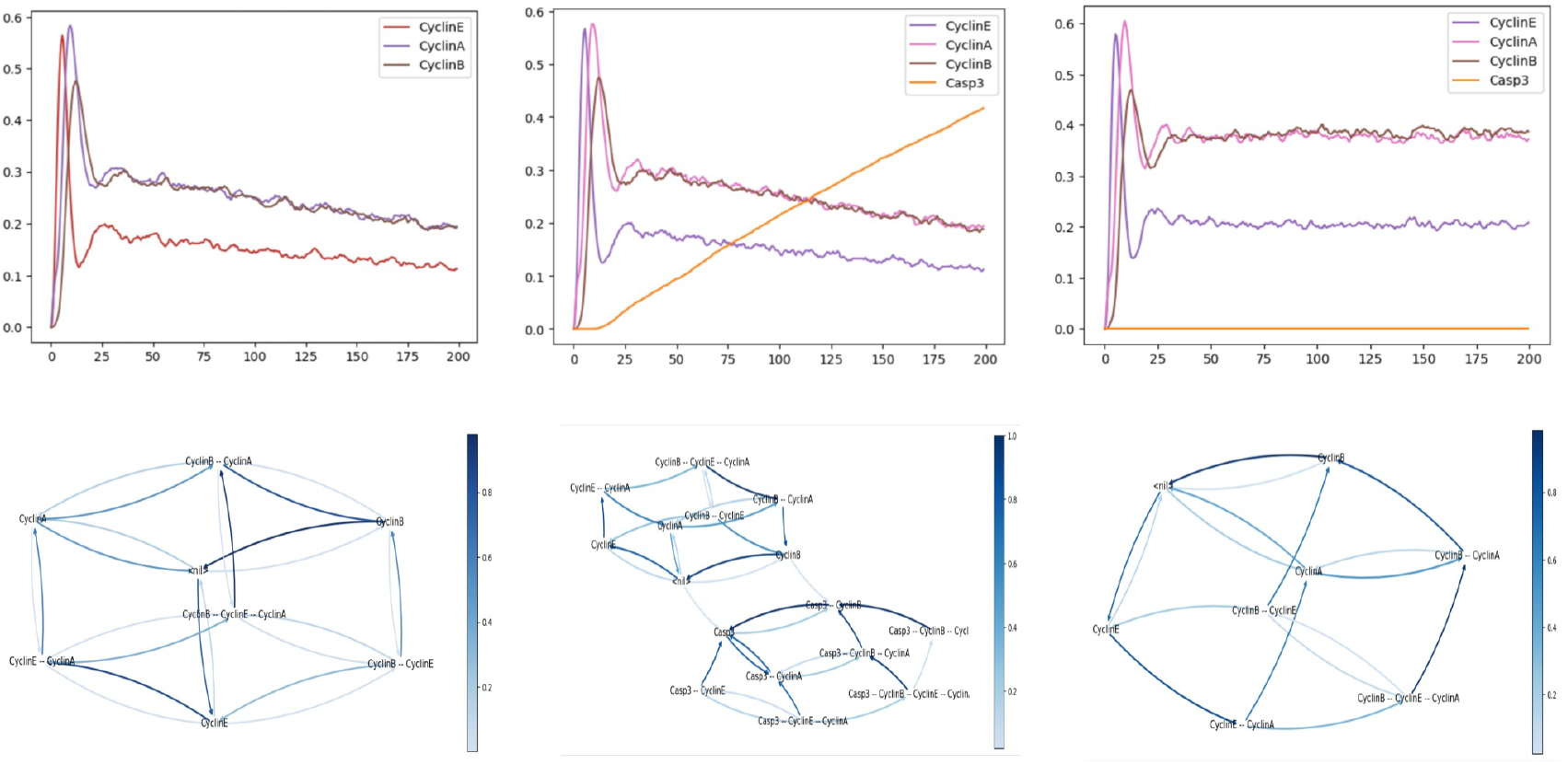
(Left) *MaBoSS* simulation of the wild-type condition, showing the cyclins activation and inactivation via their mean probabilities (Upper left panel), or via their compressed state transition graph (Lower left panel). (Middle) MaBoSS simulation of the wild-type condition, showing the cyclins and Casp 3 mean probabilities (Upper middle panel) and compressed state transition graph (Lower middle panel). (Right) MaBoSS simulation of the p110++ mutant, showing the cyclins and Casp 3 mean probabilities (Upper right panel) and compressed state transition graph (Lower right panel).

We can further compute the frequencies of cyclin activations and inactivations over all individual trajectories of the simulation. To do this, we use a new functionality of *MaBoSS* estimating the frequencies of transitions between *activity patterns* involving selected nodes and plot them as a graph. To prevent explosion of the size of this graph, we only use the three main cyclins responsible for the cyclic attractor. The following code cell computes such transition frequencies and displays the results as a *compressed state transition graph*, i.e. a projection of the transition frequencies computed over the full state space onto the cyclin activity patterns of interest.

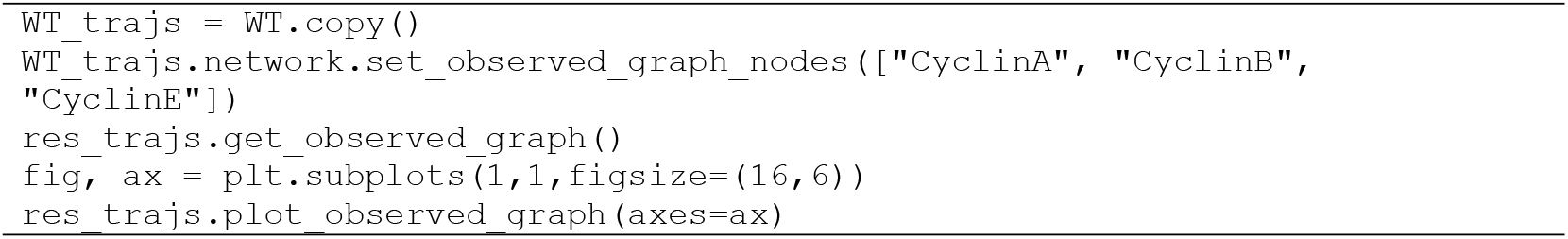

The resulting transition graph is shown in Figure 8 (bottom left).

In this graph, the transitions denoted by darker blue arrows have a higher probability of occurring. This representation enables us to visualise the most probable sequence of cyclin activations and inactivations. Starting from the state (corresponding to G0), the most probable sequence of state transitions successively involves CyclinE activation, CyclinA activation, CyclinE inactivation, CyclinB activation, CyclinA inactivation, and finally CyclinB inactivation, thereby completing the cycle. Note that other cycles can be followed with lower probabilities.

#### 3.9.2. Analysis of the apoptosis triggering timing along the cell cycle

Here we study the condition leading to apoptotic death, focusing on the point in the cell cycle where Casp3 is activated. We can achieve this using the following code cell:

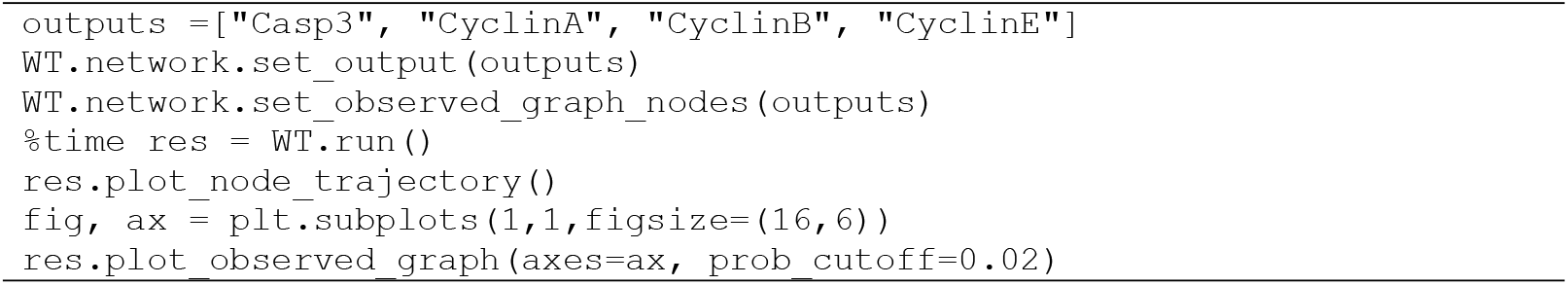

The resulting graphs are shown in **Figure 8 (middle)**. In the time plot (**Figure 8, middle top**), we can observe that cells initially follow the expected cycle, but quickly activate Casp3 and are thus led to apoptosis.

Turning to the compressed transition graph (**Figure 8, middle bottom**), we used a minimum probability threshold to avoid plotting unlikely edges. We see that there are two main ways to activate apoptosis: from <nil>, corresponding to G0, and when cyclin B is active, denoting the G2/M phase.

#### 3.9.3. Analysis of cyclin triggering timing in presence of ectopic p110_H

Finally, we can perform the same analysis in the case of a forced p110_H activation. This can be achieved using the following code cell:

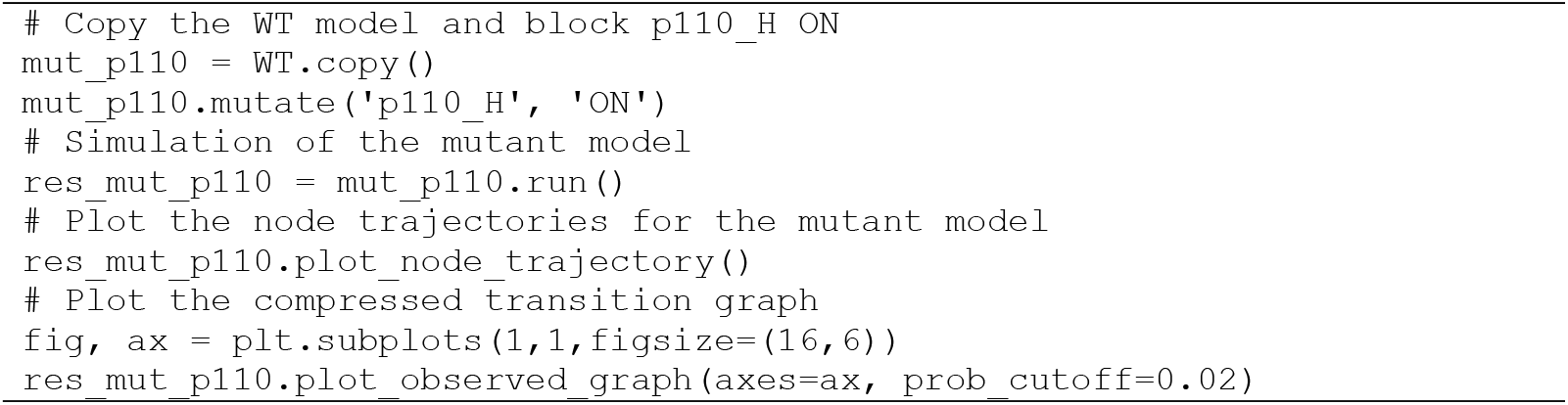

As can be seen in the resulting graph (**Figure 8, right**), the ectopic activity of p110_H completely disables the activation of apoptosis, enabling the maintenance of the oscillatory behaviour.

## 4. Conclusions

After providing the instructions to install the 20 different tools integrated in the CoLoMoTo software suite, this tutorial focused on the analysis of a published model of the regulatory network controlling mammalian cell proliferation [13].

The tutorial includes *python* code enabling the reproduction of several of the results reported by Sizek et al., as well as to further extend these results. Using selected tools included in the CoLoMoTo suite, we showed how to compute the model attractors with synchronous or asynchronous updating. Although stable states are naturally preserved, our analysis emphasises the sensitivity of synchronous cyclic attractors to update mode.

We further showed how the use of a probabilistic simulation framework, implemented in the *MaBoSS* software, can provide interesting information on the transient behaviour of such a complex model, for wild-type conditions, as well as for different types of model perturbations.

Integrating all these analyses into an executable Jupyter notebook greatly facilitates their reproducibility, as well as extensions. The notebook can also be used as a template for encoding completely new model analyses.

Due to space constraints, this notebook explicitly covers the use of only a fraction of the tools available in the CoLoMoTo collection. However, all integrated tools are accompanied by short specific tutorials available in the CoLoMoTo *Docker* container, as well as on the CoLoMoTo website.

## Acknowledgements

We are grateful to Claudine Chaouiya and Pedro Monteiro for their key contributions to the development of *GINsim*, as well as to our many colleagues who contributed tools integrated in the CoLoMoTo environment.

## Data accessibility

The notebook and required companion files are available at the url: https://github.com/colomoto/colomoto-tutorial-sizek.

The source notebook file (with a .ipynb extension) can be uploaded and executed within the Jupyter interface of the CoLoMoTo notebook, using the Docker image colomoto/colomoto-docker:2025-01- 01.

